# FtsZ is essential until the late stage of constriction

**DOI:** 10.1101/2022.03.01.482533

**Authors:** Lauren C. Corbin Goodman, Harold P. Erickson

## Abstract

There has been recent debate as to the source of constriction force during cell division. FtsZ can generate a constriction force on tubular membranes in vitro, suggesting it may generate the constriction force in vivo. However, another study showed that mutants of FtsZ did not affect the rate of constriction, whereas mutants of the PG assembly did, suggesting that PG assembly may push the constriction from the outside. Supporting this model, two groups found that cells that have initiated constriction can complete septation while the Z ring is poisoned with PC190723. PC19 arrests treadmilling but leaves FtsZ in place. We sought to determine if a fully assembled Z ring is necessary during constriction. To do this, we used a temperature sensitive FtsZ mutant, FtsZ84. FtsZ84 behaves as wild type at 30° C, but it disassembles from the Z ring within one min upon a temperature jump to 42° C. Following the temperature jump we found that cells in early constriction stop constricting. Cells that had progressed to the later stage of division finished constriction without a Z ring. These results show that in *Escherichia coli*, an assembled Z ring is essential for constriction except in the final stage, contradicting the simplest interpretation of previous studies using PC19.

## 2 Introduction

The protein FtsZ is essential to bacterial cytokinesis. Along with its membrane anchor, FtsA, it assembles a Z ring early in the cell cycle. The Z ring subsequently recruits downstream proteins that remodel the peptidoglycan (PG) wall, suggesting that it serves as a scaffold for this recruitment.

An ongoing controversy is what generates the force for constriction. In 2008, Erickson and Osawa found that Z rings containing only membrane-tethered FtsZ could constrict multilayer tubular liposomes [1]. In the presence of FtsA, rings were able to complete constriction [2]. This established the Z ring as a likely source of constriction force. The mechanism of force generation was suggested to be a transition of FtsZ protofilaments from straight to curved, generating a bending force that pulls the cell membrane inward [3].

The Z-centric constriction model recently was questioned when Coltharp and colleagues found that the rate of constriction was not altered by a mutation of FtsZ, but was slowed by a mutation of FtsI, a key component of the PG synthesis machinery [4]. They suggested that the constriction force was generated by FtsI remodeling the PG, pushing the cell membrane from the outside. Erickson and Osawa suggested an alternative explanation, that FtsZ was providing the constriction force, but the rate of constriction was limited by PG remodeling serving as a strong brake [5, 6]. However, the idea of constriction force generated by PG remodeling, with FtsZ serving primarily as a scaffold, has risen to prominence in the field.

More recently, two groups reported that constriction could continue after poisoning the Z ring with the drug PC190723 (abbreviated here as PC19). PC19 is a candidate antibiotic that targets FtsZ [7].	It causes subunits to lock into the T conformation [8] and stabilizes PFs in vitro [9]. When *Bacillus subtilis* cells were treated with to PC19, patches of FtsZ that were treadmilling around the Z ring immediately arrested their movement [10]. Experiments with PC19 in *Staphylococcus aureus* [11] and *B. subtilis* [12] showed that if the Z ring were poisoned before constriction began, PC19 blocked constriction. However, once constriction had been initiated, it would continue to complete septation in the presence of PC19.	This suggested that FtsZ treadmilling is essential for condensation and maturation of the Z ring, but is not essential for constriction. If treadmilling is not essential for constriction, perhaps FtsZ itself is dispensable once constriction has initiated.

A problem with this more global conclusion, that FtsZ may be dispensable for constriction, is that PC19 does not immediately disassemble the Z ring. PC19 and the related benzamide 8j eventually leads to disassembly of FtsZ from the Z ring and redistribution into foci scattered around the cell. However, FtsZ remains at the Z ring for 10-30 min [13].

Could the static, non-treadmilling Z ring still contribute to constriction? Importantly, FtsZ can still form both straight and curved protofilaments in the presence of PC19 [9] and 8j [13]. Thus, while PC19 rapidly arrests tread-milling, it leaves FtsZ in place and potentially able to continue generating a bending force on the membrane. This FtsZ-based force may be essential for the continued constriction after poisoning by PC19.

We sought to determine if an assembled Z ring is necessary during constriction. For this, we employed an FtsZ84 genomic mutant strain of *Escherichia coli*, JFL101, as a tool. JFL101 behaves as wild type at 30° C; however, at 42° C, the Z ring disassembles within one min. [14]. We used a rapid temperature jump (t jump) to disassemble the Z ring and observe how the cells respond to the sudden absence of FtsZ.

## 3 Methods

### 3.1 Cell cultures

All liquid cultures were grown in LB (1% tryptone, 0.5% sodium chloride, 5% yeast extract, 0.2% glucose, pH 7.0) Liquid cultures were started from −20° C freezer stock (10% glycerol) or streaked plates and grown in LB at 30° C overnight. Cultures were spun down and diluted by 200 to 1000 fold into fresh LB and grown for several hours until mid-log phase (OD_600_ between 0.15 and 0.5) for experiments.

Aliquots of 4% formaldehyde in PBS were prepared less than one month ahead of experiment. PBS was heated to 60° C, and powdered paraformaldehyde was added to a final concentration of 4%. Sodium hydroxide was titrated into the solution until the paraformaldehyde had dissolved. The solution was cooled, and the pH was adjusted to 7.0. The solution was filtered with a 0.22 μm polyethersulfone membrane filter, and aliquots were frozen at −20° C.

### 3.2 Cell counting

T-jump experiments were performed in a 37° C incubator room to reduce heat fluctuations while samples were removed and fixed. An Erlenmeyer flask with 50 mL of filtered LB was preheated to 42° C in a water bath for at least two hours prior to the start of the experiment. Pipettes, pipette tips, and 2X fixation solutions were also pre-warmed. Formaldehyde was diluted with PBS to produce 2% formaldehyde (2X fixative). The 2X fixative was filtered again within an hour of the start of the t-jump experiment. We pre-aliquoted 500 μL of 2X formaldehyde for each sample.

At the start of the t-jump experiment, mid-log cells were pre-diluted if needed into fresh 30° C LB to obtain an OD_600_ of 0.15. 1.5 mL of cells were added to the 50 mL 42° C LB. Immediately, 500 μL of cells were removed and diluted into 500 μL 2X fixative, for a final concentration of 1% formaldehyde. We consider this time = 0 min. Initially, the temperature of the LB dropped 0.3° C below 42° C, but the water bath warmed the LB to 42° C within a min. We performed this 2-fold dilution from the 42° C LB to the 2X fixation at each time point. We also recorded the LB temperature at these time points. We repeated the dilution between 3 and 5 times for each experiment at each time to check for preciseness of the single experiment. We repeated this experiment to obtain 4 replicates with our desired experimental conditions. Fixed samples were cooled to room temperature and counted within three hours.

To count the cells, we used the BD FACSCanto II flow cytometry system at medium flow rate. We counted cells for two min gated against forward scatter area and side scatter area. We collected blank measurements for each experiment by counting a solution of 500 μL fresh LB plus 500 μL 2X fixative.

We gated and counted the cells in each sample with the free and open source Python package FlowCytometryTools [15, 16]. We highly recommend this package, as it is fast with large FCS files, includes excellent documentation, and is intuitive. We performed data analysis and made figures in R with packages in the Tidyverse [17, 18].

We optimized cell concentration by performing a serial dilution and counting cells. To do this, we serially diluted the fixed cells 2 fold in filtered PBS. We counted cells for one min and plotted the cell count against the calculated cell concentration based on an ideal serial dilution.

### 3.3 Fixed membrane imaging

Fixative with 4% formaldehyde was thawed from freezer aliquots, and glutaraldehyde from frozen a 25% stock was added to a concentration of 0.4%. The fixative was filtered approximately 1 hour prior to the t jump. We premeasured 6 mL aliquots of 4X fixative and warmed it in the 37° C incubator room. We filtered and pre-warmed 100 mL of LB to 42° C.

Cells were grown to mid-log phase as previously described. At the start of the t-jump experiment, we diluted 2.5 mL of culture into 100 mL of 42° C LB. To collect a sample, we removed 20 mL of cells and combined it with 6 mL of 4X fixative. After two min, we moved the sample to ice. We repeated this for each point in our time series. For the 0 min sample, we diluted 0.5 mL of cells into 20 mL of 30° C LB and added 6 mL of 4X fixative. We stored fixed samples at 4° C overnight.

To image the fixed cells, we pelleted the cells at 4200 × g for 5 min and resuspended the cells in 1 mL LB. We pelleted the cells again and resuspended them in 100 μL fresh LB. We added FM4-64 dye (Invitrogen) to a final concentration of 0.7 μg/mL. We added 25 μL of stained cells to a 2 cm × 6 cm agar pad made with water and 1% agarose. We allowed the sample to dry for 30 min and covered it with a coverslip. We imaged the cells within 3 hours of making the pads.

We imaged the cells with STED implemented on a Leica DMi8 microscope. We excited our sample at 516 nm with a white light laser (0.7% of total power) and collected emissions between 615 nm and 685 nm with a GaAsP HyD detector. We used the 775 nm STED depletion laser at 8% power. We collected a focal series of 7 stacks over 1 μm depth. Each pixel measures 21nm × 21 nm.

We performed deconvolution with the Huygens professional software [19]. We then segmented the images with LabKit in FIJI. We used Python3 [16] and NumPy [20] to measure the width and height of each cell.

### 3.4 Live membrane imaging

Cells were grown to mid-log phase as previously described. We pelleted the cells at 1600 × g for 5 min and resuspended them in 200 μL of fresh LB. We added FM5-95 dye (Invitrogen) to a concentration of 4 μg/mL from a stock solution in water. We sonicated a glass bottom dish (MatTek) in Helmenex followed by 100 % ethanol. We plated 5 μL of cells onto the glass bottom coverslip dish and covered it with a 1% agarose LB pad, approximately 1-2 cm wide, square. We put a lid on the glass bottom dish and incubated it at 30° C for 30 min.

To perform the t jump, we transferred the glass bottom dish with the cells into a 42.5° C microscope incubator (Oko labs). We imaged the cells with a Nikon Eclipse Ti2 microscope with NIS Elements software and a heated objected. We excited the sample at 560 nm and collected emissions above 630 nm. We took a time-lapse of a single field of view with 1 frame/min.

## 4 Results

Our experiments all used JFL101 as a tool. JFL101 is a temperature sensitive FtsZ *E. coli* mutant that behaves wild type at 30° C (Fig. 1A). However, at 42° C, the Z ring disassembles within one min [14]. For our t-jump experiments, we grew JFL101 at 30° C, and diluted the cells into 42° C media, causing the Z ring to disassemble. Following this t jump we observed the process of septation with several methods.

**Figure 1:**
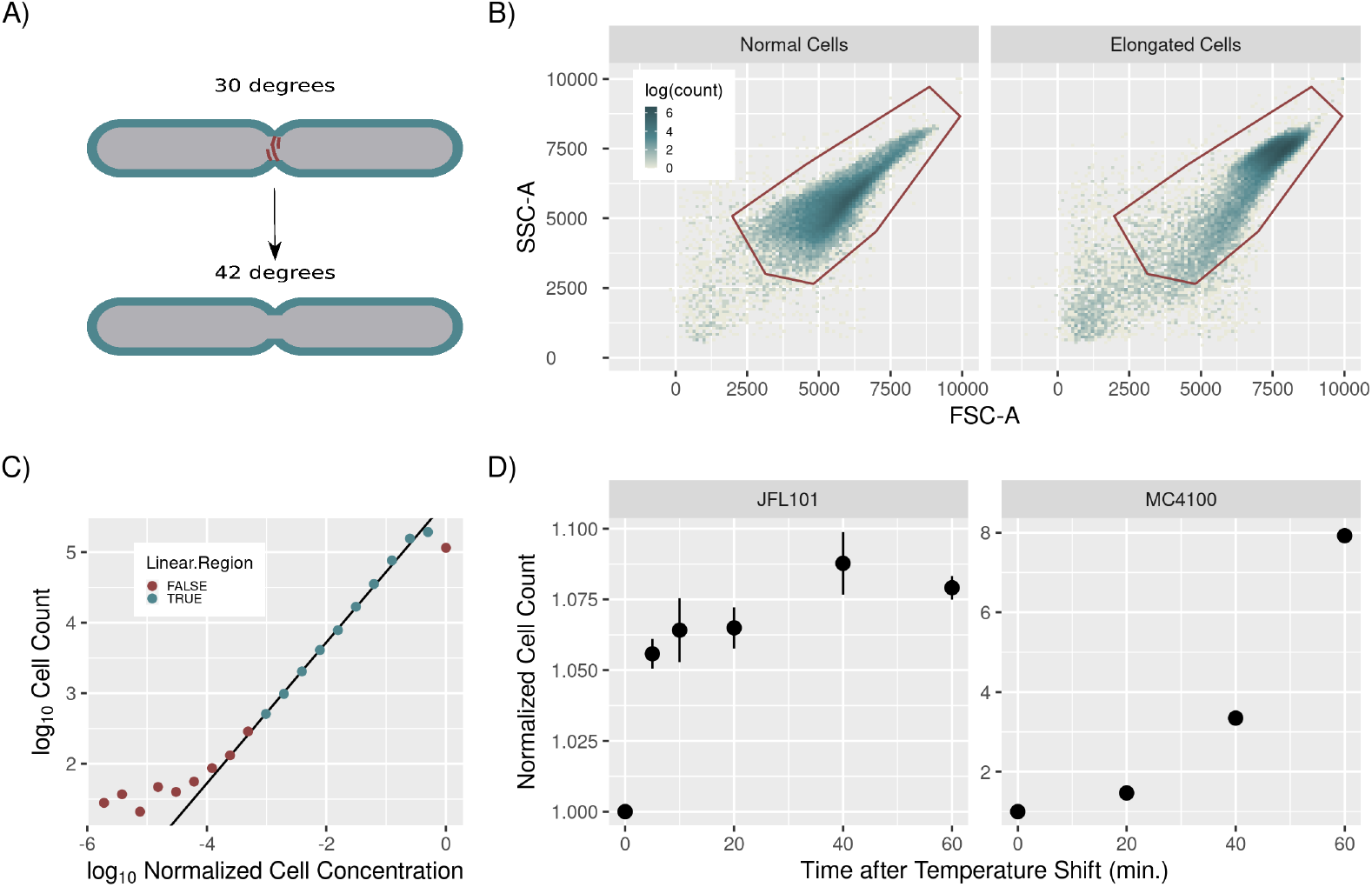
**A)** Diagram of t-jump experiment. **B)** Representative flow cytometry histograms of normal (left side) and elongated (right side) cells. Red polygons outline the signals counted as cells. SSC-A and FSC-A are log transformed side and forward scatter area. **C)** Calibrating the valid range of cell counts from a two-fold serial dilution of cells. Line represents a linear regression fit where slope = 1 and r^2^ = 0.996. **D)** Mean of normalized cell count over time after t jump of JFL101 cells (left side, n=4) or MC4100 cells (right, n=2). Note the very different y-axis scales. Cell counts for each replicate were normalized to the count at 0 min. Error bars are standard error. Note that standard error of MC4100 was too small to display with this y-axis scale.

### 4.1 Counting division by flow cytometry

Our first method characterized the t jump by counting cells to determine what fraction of cells could continue constriction to complete division following the sudden depletion of FtsZ from the Z ring. For this we fixed aliquots of cells at set time points following the t jump, and used flow cytometry to count the cells and determine a precise growth curve.

Before we began this experiment, we optimized flow cytometer settings to ensure we were capturing our cells. We were able to isolate the cells by gating against the forward and side scatter area (Fig. 1B, left panel). When JFL101 loses the Z ring, the cells elongate, so we optimized the flow rate, gate and cell density to also capture elongated cells (Fig. 1B, right panel). We found that SYTO BC bacterial stain and propidium iodide stain did not improve our ability to gate the cells.

We first determined the concentration at which we could obtain a valid count the cells. At high concentrations, cells can crowd the central fluid stream and evade being counted. At low concentration, the noise obscures the true count. For calibration, we used a serial dilution assay. After fixing a sample of cells, we serially diluted the cells two-fold and counted the cells for one min. We found the region in which a two-fold dilution would result in a two-fold reduction in cell count. This was approximately between 500 and 194,000 cells per min (Fig. 1C).

After these optimizations, we then performed the full t-jump experiment to disassemble the Z ring, collect fixed samples over time and count the cells. We normalized the cell counting results from each replicate to the replicate’s initial average cell count at 0 min. Fig. 1D, left panel, shows the combined results of four replicated t-jump experiments. When the JFL101 cells were jumped to 42° C, the cell count then increased to a plateau at 6.4-6.5% above the zero time at 10 and 20 min. From this, we conclude that 6.4% of cells can complete division without FtsZ assembled in a Z ring.

We repeated this experiment with MC4100, the wild-type *E. coli* strain from which JFL101 was constructed (Fig. 1D, right panel). We found that the t-jump treatment did not inhibit division.

### 4.2 Imaging membrane constrictions following the sudden depletion of Z ring

We then wanted to understand what happens to the constricting membrane following the sudden depletion of FtsZ. We repeated the t-jump experiment, and imaged the cells after staining the outer membrane with the fluorescent dye FM4-64 [21]. We used the super-resolution technique STED for imaging. We collected images of several hundred fixed cells at each time point and quantified the extent of constriction. These images created a pseudotime lapse of constrictions after the t jump.

Fig. 2 contains cropped images of cells sorted by their constriction progress and time. Our zero min sample was collected just before the t jump. At this time, we found a wide range of constrictions, from almost fully constricted (bottom of montage) to just beginning constriction (top of montage). 18.1% of cells were constricted at time 0 min, and this decreased to 9.0% at 15 min (table 1). The percent constricted cells decreased further at 30 min, perhaps due to relaxation of some constrictions.

**Figure 2:**
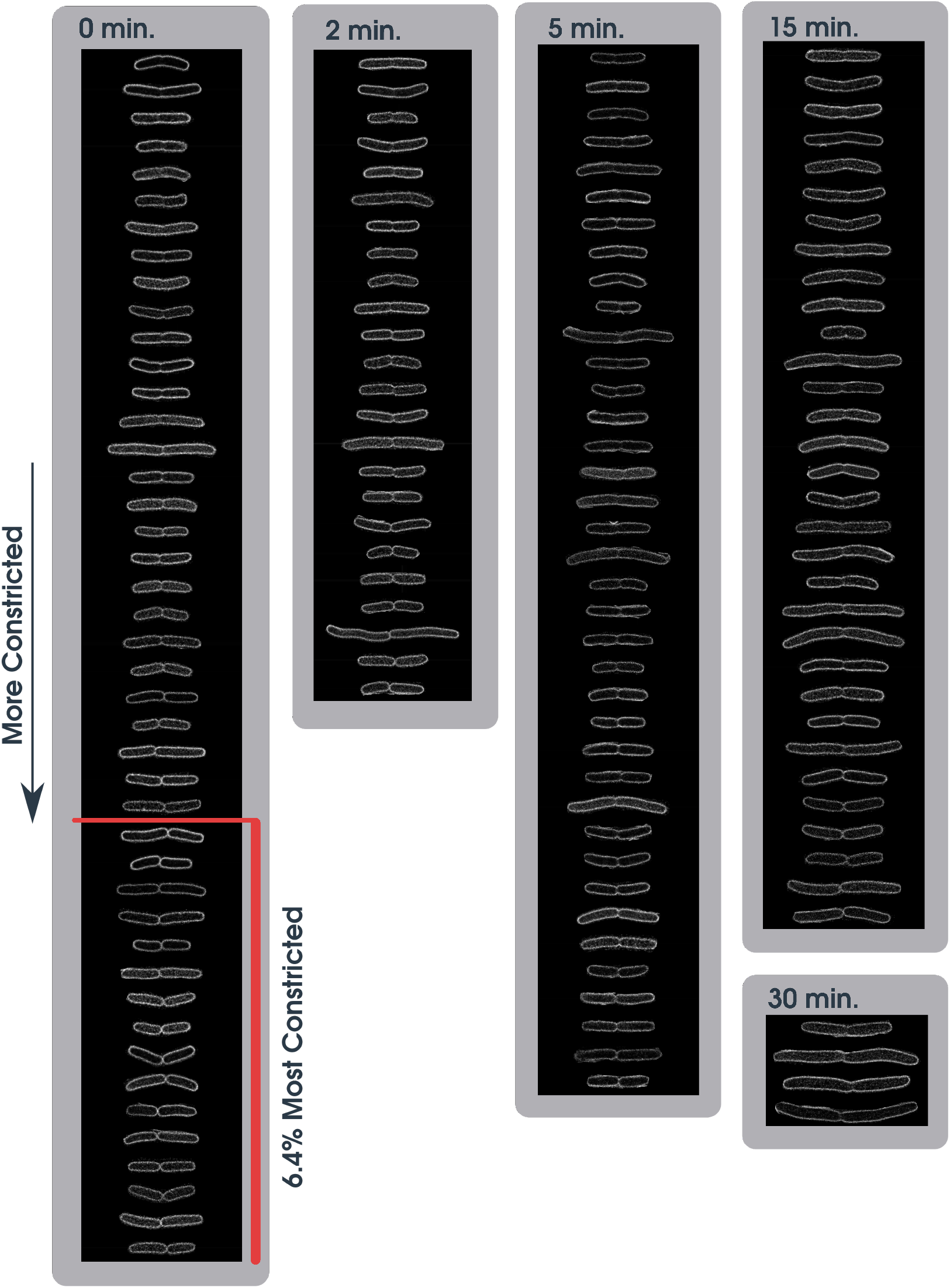
Cropped images of cells sorted by constriction progress and time point. Cells towards the bottom are the furthest constricted. For time 0 min, the red line denotes the 6.4% mo8st constricted cells (out of total cells). Approximately 80-90% of cells were not constricted and are not shown here.

**Table 1:** JFL101 cells counted and proportion with constrictions for fixed imaging analysis.

| Time (min.) | Cells Counted | Constrictions |
| --- | --- | --- |
| 0 | 371 | 18.1% |
| 2 | 175 | 14.9% |
| 5 | 318 | 14.5% |
| 15 | 379 | 9.0% |
| 30 | 103 | 4.9% |

We then measured the extent of constriction over time (Fig. 3). The normalized constriction is the constriction width divided by the average cell width. A normalized constriction near zero means that the cell has almost completed division. A normalized constriction near one means the cell has just begun constriction. The overall conclusion from Fig. 3 is that cells in early constriction are retained at 2, 5 and 15 min after the t jump, while late constricting cells disappear. The disappearance of the late constricting cells is consistent with their having completed division. To correlate these data with the cell counting, which showed a 6.4% increase in cells following the t jump, we show a line in Fig. 3 at 0.44 normalized constriction at 0 time. 6.4% of cells are below this line and presumably completed division.

**Figure 3:**
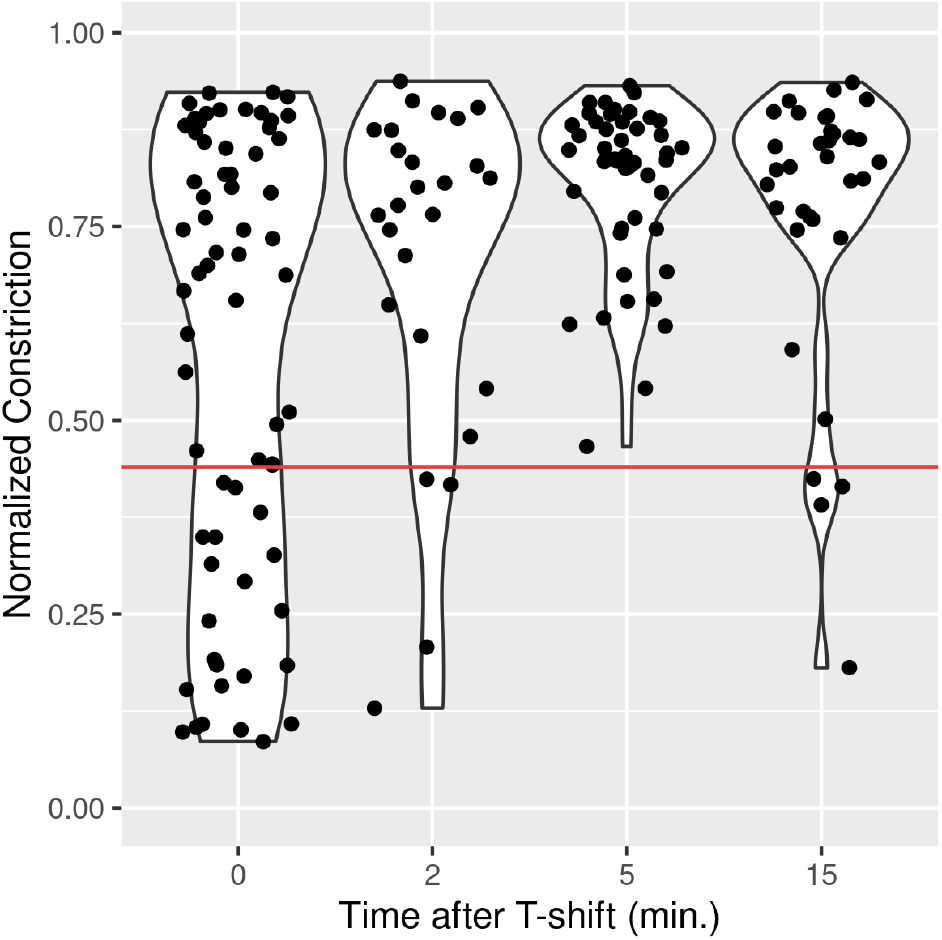
Density of normalized constriction after t jump. Unconstricted cells were omitted from this plot.

To further inform our analysis of these results, we repeated the t-jump experiment with live cells. We grew live JFL101 stained with FM5-95 on small agar pads. FM5-95 is also an outer membrane dye but was less toxic and had longer lasting fluorescence than FM4-64 in live cells. We transferred the agar pad to a 42.5° C incubator microscope and imaged the cell’s response to the t jump. We did not find any cells that initiated constriction after the t jump. If a cell was previously constricted at the start of the t jump, the cell either completed constriction (Fig. 4 B) or maintained a constant constriction (Fig. 4 A). Some cells that remained constricted appeared to slowly relax over the course of 30 min (Fig. 4 C). We were only able to image a small number of live cells, so these images are presented as examples without statistics.

**Figure 4:**
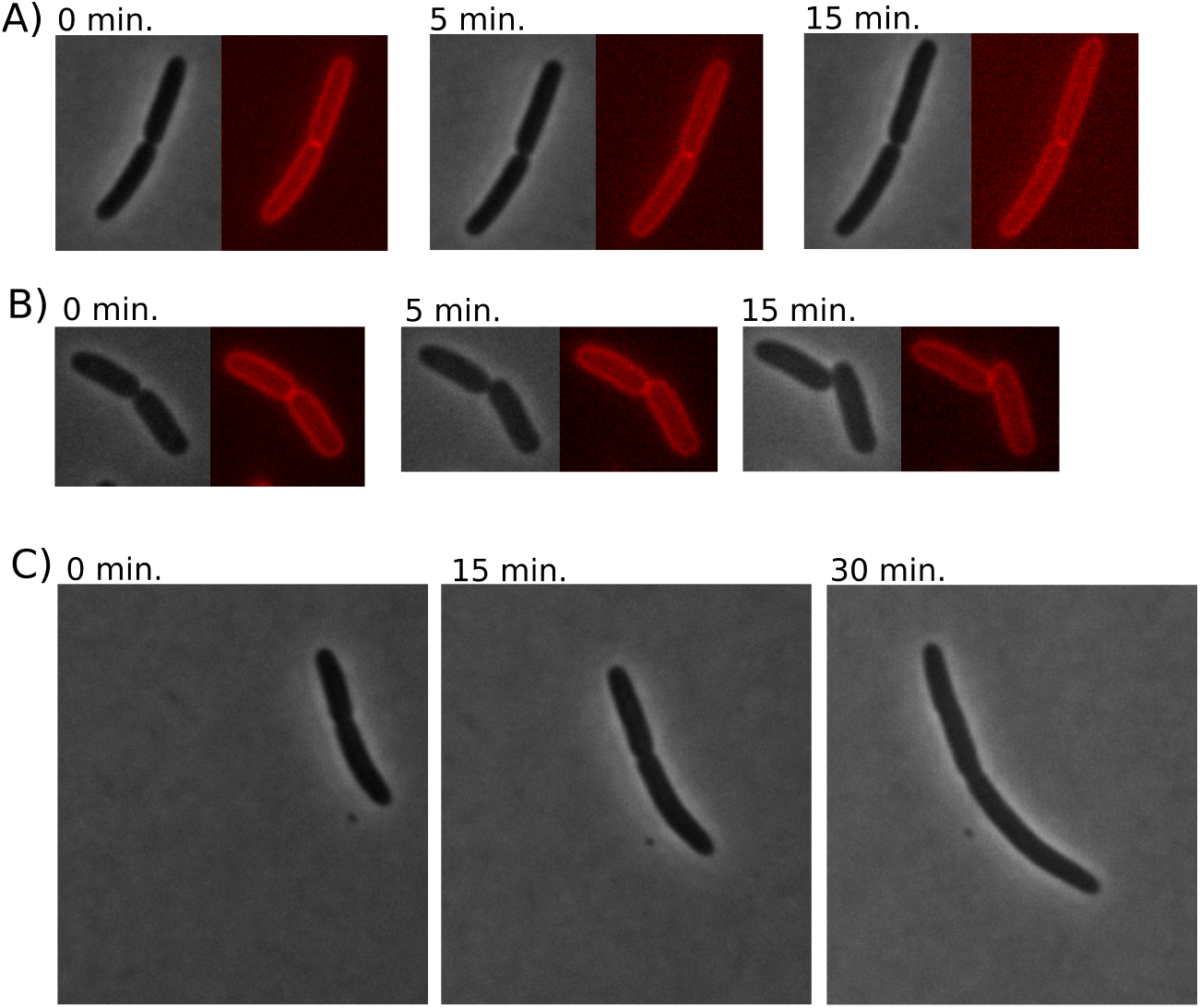
Live time lapse of JFL101 after t jump. Gray panels are phase contrast and red panels are FM5-95 dye. Cells either maintained constriction (**A**), completed constriction (**B**), or slowly relaxed (**C**).

## 5 Discussion

Two previous studies used the compound PC19 to poison the Z ring in *S. aureus* [11] and *B. subtilis* [12]. Both studies found that PC19 blocked initiation of new constrictions, but constrictions that had already begun continued to complete septation. Since PC19 rapidly arrests treadmilling, these results suggest that treadmilling is needed for formation and maturation of the Z ring and to initiate constriction, but treadmilling was not needed for constriction itself. But since PC19 leaves FtsZ in the Z ring for 10-30 min [13], it is possible that the static FtsZ was still contributing to constriction.

The temperature-sensitive FtsZ84 provides a way to rapidly and completely remove FtsZ from the divisome. Addinall et al. observed that cells retained sharp constrictions two min after the T jump, and constrictions remained but became more blunt or rounded after ten min [14]. This is different from the case of *S. aureus* and *B. subtilis* following PC19 poisoning, where all constrictions disappeared as they completed septation. However, the observations of Addinall et al. were based on scanning EM images, and constricted cells were not counted quantitatively.

Our study provides quantitation. Following the t jump, we found a 6.4% increase in cell count at 10-15 min (Fig. 1D, left panel). Since 18.1% of cells showed a constriction at 0 min, this suggests that about one third of the constricted cells completed division. Light microscopy showed that cells in late constriction disappeared at 2-15 min, while the fraction of cells in early constriction remained about the same. Apparently cells in early constriction become locked and unable to progress when FtsZ is depleted. This is different from the results with PC19 poisoning, where even cells in early constriction progressed to complete septation [12, 11].

That the late constricted cells can complete division without FtsZ is not unexpected. Söderström et al. have shown that FtsZ disassembles from the Z ring before constriction is complete [22, 23]. In Söderstöm et al. 2016, the last two frames in their Fig. 5b show FtsZ still present at a constriction width of 0.3 μm and gone at a width of 0.2 μm. This shows that cells naturally complete constriction without FtsZ from 0.3 – 0.2 μm. Our work suggests that they may be able to do so from a somewhat earlier point if FtsZ is artificially depleted. We have drawn the line in Fig. 3 at 0.44 normalized constriction, to show the 6.4% of cells completing division in our cell counting measurement.

**Figure 5:**
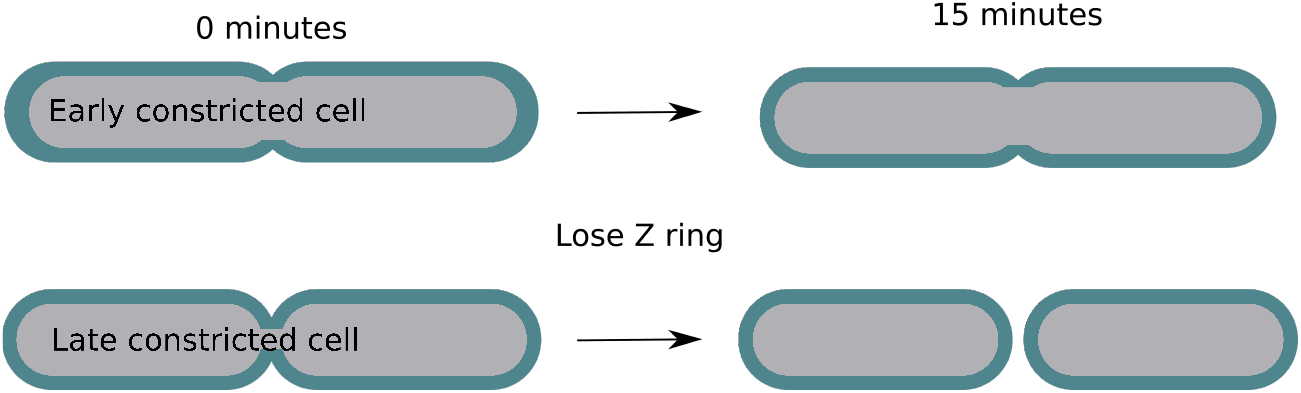
Diagram of early vs. late cells at zero and 15 min from our experiments.

A potentially important difference in our study and those of Monteiro et al. [11] and Whitely et al. [12] is the bacterial species. *S. aureus* and *B. subtilis* are gram-positive bacteria while our study was of gram-negative *E. coli*. In gram-positive bacteria, constriction involves a sharp septum at all stages, while gram-negative bacteria have a shallower V-shaped constriction at earlier stages [24, 25] (but see [5] for a hypothesis uniting the two mechanisms). It will therefore be important to determine if gram-positive bacteria require FtsZ for early constriction.

What generates the force for the late constriction? One possibility is PG synthesis pushing the membrane from the outside [4]. This may require a Brownian ratchet mechanism to move the membrane from contact with PG remodeling enzymes, in order to insert new glycan strands. Such a mechanism has not been developed, and may encounter problems [26].

A third potential force generator is excess membrane production. Once a membrane furrow has been initiated at the division site, excess membrane would push this furrow to continue constriction, rather than initiate new furrows [6]. Evidence for this has come from experiments with L form bacteria, which are bacteria that are modified to survive without their PG cell wall. Leaver and colleagues found that L form *B. subtilis* do not require FtsZ to divide [27]. Instead, it divides with blebs created by excess membrane [28]. That study proposed that excess membrane production may have been the driving force for division of the earliest life form. An *E. coli* L form has been engineered to switch back to the walled state, and it can still divide without FtsZ, although it had switched from the normal cylindrical shape to a branched and bulging, ramified shape, called ‘coli-flower’ [29]. Excess membrane production could therefore serve as a primary force for late constriction in the normal bacterial cells, with PG synthesis following and filling the gap produced by the advancing membrane.

## 6 Conclusions

In contrast to the results of Monteiro et al. [11] and Whitley et al. [12], where most or all cells that had initiated constriction could complete septation after poisoning with PC19, we found that cells in late constriction could continue after depletion of FtsZ, but cells in early constriction were blocked. There are two important differences in these studies. Those of Monteiro and Whitley used gram-positive bacteria, whereas we used gram-negative *E. coli*. Therefore, the results may be species dependent. Also, their studies poisoned treadmilling but left the FtsZ assembled in the Z ring, whereas our system rapidly disassembled FtsZ from the Z ring. We think the depletion of FtsZ is likely the most important reason that early constriction was arrested in our experiments, but this would need to be addressed by similar experiments in gram-positive bacteria.

## 7 Acknowledgements

We would like to thank Alex Bisson for the generous use of his lab for live imaging and guidance on the subject. Also, we would like to thank Theopi Rados for teaching us the live imaging techniques and helping develop our experimental protocol.

## References

[1] M. Osawa, D. E. Anderson, and H. P. Erickson, “Reconstitution of contractile ftsz rings in liposomes,” Science, vol. 320, no. 5877, pp. 792–4, 2008.

[2] M. Osawa and H. P. Erickson, “Liposome division by a simple bacterial division machinery,” Proc Natl Acad Sci U S A, vol. 110, no. 27, pp. 11000–4, 2013.

[3] H. P. Erickson, D. E. Anderson, and M. Osawa, “Ftsz in bacterial cytokinesis: Cytoskeleton and force generator all in one,” Microbiology and Molecular Biology Reviews, vol. 74, no. 4, pp. 504–528, 2010.

[4] C. Coltharp, J. Buss, T. M. Plumer, and J. Xiao, “Defining the rate-limiting processes of bacterial cytokinesis,” Proc Natl Acad Sci U S A, 2016.

[5] H. P. Erickson, “How bacterial cell division might cheat turgor pressure - a unified mechanism of septal division in gram-positive and gram-negative bacteria,” BioEssays, vol. 39, no. 8, p. 1700045, 2017.

[6] M. Osawa and H. P. Erickson, “Turgor pressure and possible constriction mechanisms in bacterial division,” Frontiers in Microbiology, vol. 9, p. 111, 2018.

[7] D. J. Haydon, N. R. Stokes, R. Ure, G. Galbraith, J. M. Bennett, D. R. Brown, P. J. Baker, V. V. Barynin, D. W. Rice, S. E. Sedelnikova, J. R. Heal, J. M. Sheridan, S. T. Aiwale, P. K. Chauhan, A. Srivastava, A. Taneja, I. Collins, J. Errington, and L. G. Czaplewski, “An inhibitor of ftsz with potent and selective anti-staphylococcal activity,” Science, vol. 321, no. 5896, pp. 1673–5, 2008.

[8] T. Matsui, J. Yamane, N. Mogi, H. Yamaguchi, H. Takemoto, M. Yao, and I. Tanaka, “Structural reorganization of the bacterial cell-division protein ftsz from staphylococcus aureus,” Acta Crystallogr., Sect. D: Biol. Crystallogr., vol. 68, no. Pt 9, pp. 1175–88, 2012.

[9] J. M. Andreu, C. Schaffner-Barbero, S. Huecas, D. Alonso, M. L. Lopez-Rodriguez, L. B. Ruiz-Avila, R. Nunez-Ramirez, O. Llorca, and A. J. Martin-Galiano, “The antibacterial cell division inhibitor pc19723 is an ftsz polymer-stabilizing agent that induces filament assembly and condensation,” J Biol Chem, vol. 285, no. 19, pp. 14239–46, 2010.

[10] A. W. Bisson-Filho, Y. P. Hsu, G. R. Squyres, E. Kuru, F. Wu, C. Jukes, Y. Sun, C. Dekker, S. Holden, M. S. VanNieuwenhze, Y. V. Brun, and E. C. Garner, “Treadmilling by ftsz filaments drives peptidoglycan synthesis and bacterial cell division,” Science, vol. 355, no. 6326, pp. 739–743, 2017.

[11] J. M. Monteiro, A. R. Pereira, N. T. Reichmann, B. M. Saraiva, P. B. Fernandes, H. Veiga, A. C. Tavares, M. Santos, M. T. Ferreira, V. Macário, and et al., “Peptidoglycan synthesis drives an ftsztreadmilling-independent step of cytokinesis,” Nature, vol. 554, no. 7693, pp. 528–532, 2018.

[12] K. D. Whitley, C. Jukes, N. Tregidgo, E. Karinou, P. Almada, Y. Cesbron, R. Henriques, C. Dekker, and S. Holden, “Ftsz treadmilling is essential for z-ring condensation and septal constriction initiation in bacillus subtilis cell division,” Nature Communications, vol. 12, p. 2448, 2021.

[13] D. W. Adams, L. J. Wu, L. G. Czaplewski, and J. Errington, “Multiple effects of benzamide antibiotics on ftsz function,” Molecular Microbiology, vol. 80, no. 1, pp. 68–84, 2011.

[14] S. G. Addinall, C. Cao, and J. Lutkenhaus, “Temperature shift experiments with an ftsz84(ts) strain reveal rapid dynamics of ftsz localization and indicate that the z ring is required throughout septation and cannot reoccupy division sites once constriction has initiated.,” Journal of bacteriology, vol. 179, no. 13, pp. 4277–4284, 1997.

[15] Jonathan Friedman and Eugene Yurtsev, “Flowcytometrytools.”

[16] Guido van Rossum, “Python3.”

[17] R Core Team, R: A Language and Environment for Statistical Computing. R Foundation for Statistical Computing, Vienna, Austria, 2021.

[18] H. Wickham, M. Averick, J. Bryan, W. Chang, L. D. McGowan, R. François, G. Grolemund, A. Hayes, L. Henry, J. Hester, M. Kuhn, T. L. Pedersen, E. Miller, S. M. Bache, K. Müller, J. Ooms, D. Robinson, D. P. Seidel, V. Spinu, K. Takahashi, D. Vaughan, C. Wilke, K. Woo, and H. Yutani, “Welcome to the tidyverse,” Journal of Open Source Software, vol. 4, no. 43, p. 1686, 2019.

[19] S. V. Imaging, “Huygens professional.”

[20] C. R. Harris, K. J. Millman, S. J. van der Walt, R. Gommers, P. Virtanen, D. Cournapeau, E. Wieser, J. Taylor, S. Berg, N. J. Smith, R. Kern, M. Picus, S. Hoyer, M. H. van Kerkwijk, M. Brett, A. Haldane, J. Fernández del Río, M. Wiebe, P. Peterson, P. Gérard-Marchant, K. Sheppard, T. Reddy, W. Weckesser, H. Abbasi, C. Gohlke, and T. E. Oliphant, “Array programming with NumPy,” Nature, vol. 585, pp. 357–362, 2020.

[21] T. Pilizota and J. W. Shaevitz, “Plasmolysis and cell shape depend on solute outer-membrane permeability during hyperosmotic shock in e. coli,” Biophysical Journal, vol. 104, no. 12, pp. 2733–2742, 2013.

[22] B. Söderström, K. Skoog, H. Blom, D. S. Weiss, G. von Heijne, and D. O. Daley, “Disassembly of the divisome in escherichia coli: evidence that ftsz dissociates before compartmentalization,” Molecular Microbiology, vol. 92, no. 1, pp. 1–9, 2014.

[23] B. Söderström, K. Mirzadeh, S. Toddo, G. von Heijne, U. Skoglund, and D. O. Daley, “Coordinated disassembly of the divisome complex in escherichia coli,” Mol Microbiol, vol. 101, no. 3, pp. 425–38, 2016.

[24] P. Szwedziak, Q. Wang, T. A. Bharat, M. Tsim, and J. Lowe, “Architecture of the ring formed by the tubulin homologue ftsz in bacterial cell division,” Elife, vol. 3, p. e04601, 2014.

[25] E. M. Judd, L. R. Comolli, J. C. Chen, K. H. Downing, W. E. Moerner, and H. H. McAdams, “Distinct constrictive processes, separated in time and space, divide caulobacter inner and outer membranes,” Journal of Bacteriology, vol. 187, no. 20, pp. 6874–6882, 2005.

[26] T. D. Pollard, “Theory from the oster laboratory leaps ahead of experiment in understanding actin-based cellular motility,” Biophysical Journal, vol. 111, pp. 1589–1592, Oct 2016.

[27] M. Leaver, P. Domínguez-Cuevas, J. M. Coxhead, R. A. Daniel, and J. Errington, “Life without a wall or division machine in bacillus subtilis,” Nature, vol. 457, no. 7231, pp. 849–853, 2009.

[28] R. Mercier, Y. Kawai, and J. Errington, “Excess membrane synthesis drives a primitive mode of cell proliferation,” Cell, vol. 152, no. 5, pp. 997–1007, 2013.

[29] R. Mercier, Y. Kawai, and J. Errington, “Wall proficient e. coli capable of sustained growth in the absence of the z-ring division machine,” Nature Microbiology, vol. 1, no. 8, p. 16091, 2016.

